# Bioinformatics analysis quantifies neighborhood preferences of cancer cells in Hodgkin lymphoma

**DOI:** 10.1101/228981

**Authors:** Jennifer Scheidel, Hendrik Schäefer, Jöerg Ackermann, Marie Hebel, Tim Schäfer, Claudia Döering, Sylvia Hartmann, Martin-Leo Hansmann, Ina Koch

## Abstract

**Motivation:** Hodgkin lymphoma is a tumor of the lymphatic system and represents one of the most frequent lymphoma in the Western world. It is characterized by Hodgkin cells and Reed-Sternberg cells, which exhibit a broad morphological spectrum. The cells are visualized by immunohistochemical staining of tissue sections. In pathology, tissue images are mainly manually evaluated, relying on the expertise and experience of pathologists. Computational quantification methods become more and more essential to evaluate tissue images. In particular, the distribution of cancer cells is of great interest.

**Results:** Here, we systematically quantified and investigated cancer cell properties and their spatial neighborhood relations by applying statistical analyses to whole slide images of Hodgkin lymphoma and lymphadenitis, which describes a non-cancerous inflammation of the lymph node. We differentiated cells by their morphology and studied the spatial neighborhood relation of more than 400,000 immunohistochemically stained cells. We found that, according to their morphological features, the cells exhibited significant preferences for and aversions to cells of specific profiles as nearest neighbor. We quantified differences between Hodgkin lymphoma and lymphadenitis concerning the neighborhood relations of cells and the sizes of cells. The approach can easily be applied to other cancer types.

## 1 Introduction

Hodgkin lymphoma (HL) is a tumor of the lymphatic system, which originates from B-lineage cells at various stages of development [18]. The annual incidence of about three cases per 100, 000 persons makes HL to one of the most frequent lymphomas of the Western civilization [16, 17]. The World Health Organization categorizes HL into different subtypes [12], mainly the classical HL (cHL) and the nodular lymphocyte-predominant HL. The cHL is further mainly divided into the nodular sclerosis cHL (NScHL) and the mixed cellularity cHL (MCcHL). About 95% of all HL are diagnosed as cHL, which is characterized by the occurrence of morphologically huge pleomorphic tumor cells called Hodgkin and Reed-Sternberg (HRS) cells. Whereas Hodgkin cells are mononucleated, Reed-Sternberg cells are multinucleated (Figure S1, supplementary material). HRS cells exhibit a broad morphological spectrum. In contrast to other cancer types, for cHL, only 1 to 2% of the tissue of the lymph node consists of tumor cells.

For diagnosis, tissue sections are cut, and the HRS cells are visualized using immunohistochemical staining with CD30. During the manual, visual inspection, pathologists pay attention to specific patterns of tumor cells in the lymph node tissue. Based on their knowledge and experience, pathologists are able to diagnose the cancer type and to prognose the disease progression. A systematic computational analysis is not yet regularly used in these processes, but would additionally provide valuable, quantified information [20].

Digital pathology is an approaching field and becomes more and more important since whole slide scanning devices allow to digitize whole tissue sections. Various imaging approaches have been applied to analyze and classify malignant tissues, see, e.g., [13, 7]. These approaches are mainly based on color, texture, and object descriptors. Morphology descriptors have been applied to describe and differentiate cell types [22] and to categorize cell nuclei [14]. Malignant cells have been reported to develop abnormal, irregularly shaped nuclei [5, 27]. Morphologic features have been used to separate and label benign and malignant tissues [15].

The analysis of morphologic cell features is important to understand the tumor development and supports computer-aided diagnosis of malignant diseases. Novkovic *et al.* have applied a graph theory-based approach to determine topological properties and robustness of the network of fibroblastic reticular cells in lymph nodes in mice. They have demonstrated the high topological robustness of the network and the critical role of network integrity for the activation of adaptive immune responses [21]. Huang et *al.* have developed a platform for multi-scale analysis, using, among others, graphics processing unit (GPU) technologies to accelerate processing of whole slide images (WSI). They have implemented their methods within a computer-aided breast biopsy analysis based on histopathological images [11].

It is so far not possible to perform life imaging of human lymph nodes affected by HL, and a corresponding mouse model of HL does not exist. The aim of the present study was to draw conclusions about HRS cell dissemination by analyzing the distribution of HRS cells in the tissue as a function of their size and shape. Do HRS cells come in close spatial contact to communicate and cooperate with each other? For example, one hypothesis was that HRS cells of elongated shape are moving, whereas frayed cells are more communicating with cells of other types. To the best of our knowledge, these questions have not been addressed so far.

There were two aspects to motivate the work. The first aspect concerns the exploration of migration of HRS cells to understand the progression of cHL. In particular, we wanted to systematically characterize and quantify the distribution of CD30-positive cells in the lymph node. Since histological sections represent snapshots of the tumor development, we explored features like the number, distance, and neighborhood relationship of tumor cells by applying statistical methods. The second aspect concerns the computational methods for the analysis of WSI including topological properties.

In this study, we considered 35 WSI of tissue sections of the cHL subtypes NScHL and MCcHL, as well as images of an inflammation of the lymph node called lymphadenitis (LA) with and without follicular hyperplasia. Based on morphological features of the cells, such as eccentricity, solidity, and area, we defined profiles classes (PC) of CD30–positive cells. We classified the cells into eight PC by specification of thresholds to distinguish between morphological categories, small and large, round and elongated, and frayed and not frayed, respectively. We determined the fraction of cells of a given PC, analyzed the distribution of cell profile diameters, and compared the results for the different diagnoses.

To address the question whether cells of a given morphological PC prefer or avoid the neighborhood of a specific PC, we studied the nearest neighborhood relationship of cell profiles. The aim was to detect statistically significant correlations. We determined the distribution of distances to the nearest neighbor to check whether preferred neighborhood relations were based on attractions between cells or on the different motility of the cells in the tissue.

## 2 Methods

### Images

We analyzed 35 two-dimensional histological WSI of human lymph nodes provided by the Dr. Senckenberg Institute of Pathology, Frankfurt am Main. The images cover the three medical diagnoses: NScHL (12 images), MCcHL (12 images), and LA (11 images). They are CD30-immunohistochemically stained to visualize the HRS cells and activated B and T lymphocytes [10]. We considered more than 400,000 CD30–positive cells in images with resolution of 0.25 *μ*m per pixel.

### Cell detection and classification

We identified cell profiles of CD30–positive cells by applying an in-house software pipeline [26, 25]. The pipeline neglected small objects of size 109 *μπ*^2^ or less. The size threshold corresponds to the size of a round cell with a diameter of 11.8 *μ*m. Applying the shape descriptors, eccentricity, solidity, and area provided by CellProfiler [3], we assigned each cell profile to one of the eight classes, see Figure 1. The eccentricity measures the deviation of a fitted ellipse from a circle. Solidity is the fraction of the object pixels in the convex hull of the cell. Together with pathologists, we defined empirically specific thresholds to distinguish between small and large, round and elongated, and frayed and not frayed cell profiles. Table S1 in the supplementary material lists the thresholds of the class definitions.

**Figure 1:**
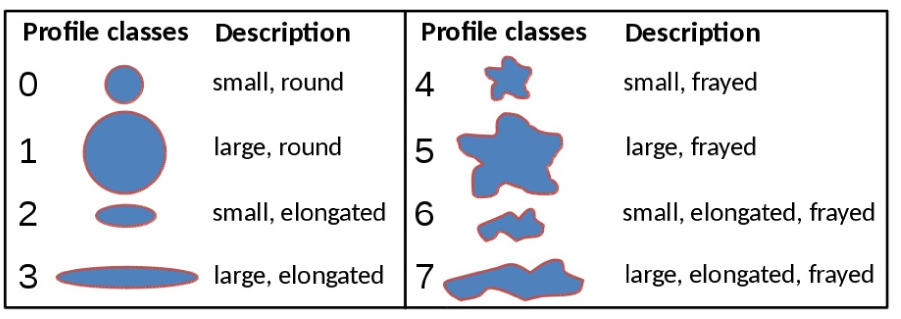
The eight profile classes (PC) and their morphological descriptions.

For each cell profile, we computed the maximal Feret diameter using the CellProfiler module *MeasureObjectSizeShape.* The Feret diameter describes the distance of two parallel tangents to the cell in a given orientation. The maximal Feret diameter referred in the text as diameter, is the largest value of all possible orientations.

### Neighborhood analysis

To detect statistically significant correlations between cell profiles, we studied the nearest neighbor relationships defined by the Euclidean distance between the centers of gravity of cell profiles. We allowed a maximal distance of 175 *μ*m (700 pixels) of a cell to its nearest neighbor. Cells without any cell within the maximal distance were called itisolated and were neglected in the neighborhood analysis. The threshold value of 175 *μ*m corresponds to ten times the diameter of an average cell and is sufficiently small to enable intercellular communication based on, e.g., chemokines or cytokines. We computed a neighborhood list, which contained for each cell the PC and the itneighbor profile class (NPC). The number of rows that has the entry, PC = *i*, determined the relative frequency, *f*(*i*), of a PC *i* ∈ {0,…,7}. Figure S2 in the supplementary material depicts an exemplary sub-section of a histological image and the corresponding neighborhood list. For a given image (ID = 1721), part (A) of Table S2 in the supplementary material exemplifies the measured probability *P*(PC = *i*) ≡ *f*(*i*) to find a cell of a certain PC.

**Table 1:** Significance matrix for the image ID = 1721 of the diagnosis MCcHL. PC stands for profile class and NPC for neighbor profile class. The blank entries stand for ns - none significant, sh for significantly high, and sl for significantly low.

| PC | 0 | 1 | 2 | 3 | 4 | 5 | 6 | 7 |
| --- | --- | --- | --- | --- | --- | --- | --- | --- |
| NPC |  |  |  |  |  |  |  |  |
| 0 | sh |  |  |  |  | sl | sl |  |
| 1 |  | sh | sh | sh |  | sh |  | sh |
| 2 |  |  | sh | sh | sl |  |  |  |
| 3 |  | sh |  | sh | sl | sh |  | sh |
| 4 |  | sl | sl | sl | sh | sl |  | sl |
| 5 |  | sh |  | sh | sl | sh |  | sh |
| 6 | sl |  |  |  |  |  |  |  |
| 7 | sl | sh |  | sh | sl | sh |  | sh |

### Statistical significance

The probability to randomly choose a neighbor of a PC, *NPC* = *j*, is proportional to the frequency of the PC, *f*(*PC* = *j*), in the image. In a random choice, the PC of the cell itself should have no influence on the selection of the nearest neighbor. We expect to measure a value for the conditional probability that is within statistical fluctuation indistinguishable from the relative frequency of the NPC, i.e., *P*(NPC = *j* | PC = *i*) ≈ *P*(PC = *j*). For example, the entry for the conditional probability, *P*(NPC = 0 | PC = 0) = 0.354, in Table S2 (B) (supplementary material) has to be compared with the entry for the expected probability, *P*(PC = 0) = 0.316, in Table S2 (A) (supplementary material).

We evaluated whether the deviation of the conditional probability from the expected probability could be a statistically significant justification for a rejection of the null hypothesis of a random selection of the nearest neighbor. For a set of *n* cells of PC = *i*, the probability to have a subset of *k* cells with nearest neighbor of NPC = *j* is given by the binomial distribution

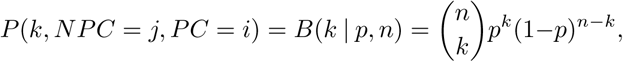

where *p* = *P*(PC = *j*) is the probability to find a class, *j*, by chance, i.e., *f*(*j*), the relative frequency of class *j*. For each image and each combination of morphological classes (PC = *i*, NPC = *j*) with *i*,*j* ∈ {0,1,…,7}, we computed the lower and upper endpoint of the prediction interval by

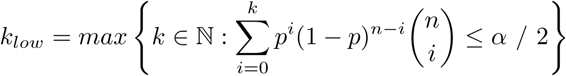

and

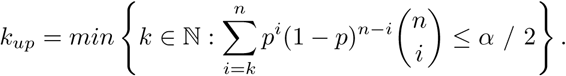

We chose a significance level of *α* = 1%. A measured value *k*, has a p-value smaller than *α* / 2 only if *k* ∉ [*k_iow_*,*k_up_*]. A *k* outside the prediction interval has a significance level of *α* = 1% which is sufficiently high enough to reject the null hypothesis of a random selection of the nearest neighbor.

For example, the image with ID = 1721 of diagnosis MCcHL contains 3435 cells of PC = 0 from 10, 860 cells in total. We computed the prediction interval [1047,1125] for the significance level *α* = 1%. We measured a number of *k* = 1217 pairs of PC = 0 and NPC = 0, which is above the upper endpoint of the 1% prediction interval. Thus, we can reject the null hypothesis for the significance level *α* = 1%. That means, within a significance smaller than 1%, the small, round cells are enriched in the neighborhood of other small, round cells. The small, round cells prefer the neighborhood of their own kind. We call such a preferred neighborhood relation to be *significantly high (sh)*.

If the number of measured pairs is smaller than the prediction interval for the significance level, we call the neighborhood relation to be *significantly low (sl)*. A *none significant (ns)* neighborhood relation denotes a number of pairs within the prediction interval. We compiled the results for each image in a significance matrix, see, e.g., Table 1. We computed an individual significance matrix for each of the 12 images of diagnosis NScHL, 12 images of diagnosis MCcHL and 11 images of diagnosis LA. The significance matrix in Table 1 exemplifies cell preferences and aversions for image ID = 1721 of diagnosis MCcHL. For example, the row for PC = 0 contains one entry itsh and two entries itsl. The entries indicate that, the class PC = 0 prefers other cells of the same class, NPC = 0, but rejects cells of NPC = 6 and NPC = 7. The empty entries in the row for PC = 0 indicated that, no preferences or aversion were measured for the combination of PC = 0 with NPC = 1, 2, 3, 4, or 5.

For each of the 64 combinations of morphological classes, we counted the number of images, in which the combination was *sh* and *sl*, respectively. The difference of these numbers gave an integer score of neighborhood relation for each combination of classes and each diagnosis, see Figure S3 in the supplementary material. The score of neighborhood relation gives a high positive value when a combination of classes is *sh* in the majority of the images, and reversely, a high negative value when a combination of classes is *sl* in the majority of the images.

## 3 Results

### Small cells occur much more frequently than large cells

Figure 2 shows the fraction of PC averaged over the 35 images. PC = 0 is the most frequently occurring class with 38.63%, describing small, round cells, followed by PC = 4 with 25.47%, standing for also small, but frayed cells. Only very few cell profiles, 0.75%, were large and elongated (PC = 3). Overall, PC = 1, 3, 5, and 7, describing large cells, were less frequent than the PC = 0, 2, 4, and 6, describing small cells.

**Figure 2:**
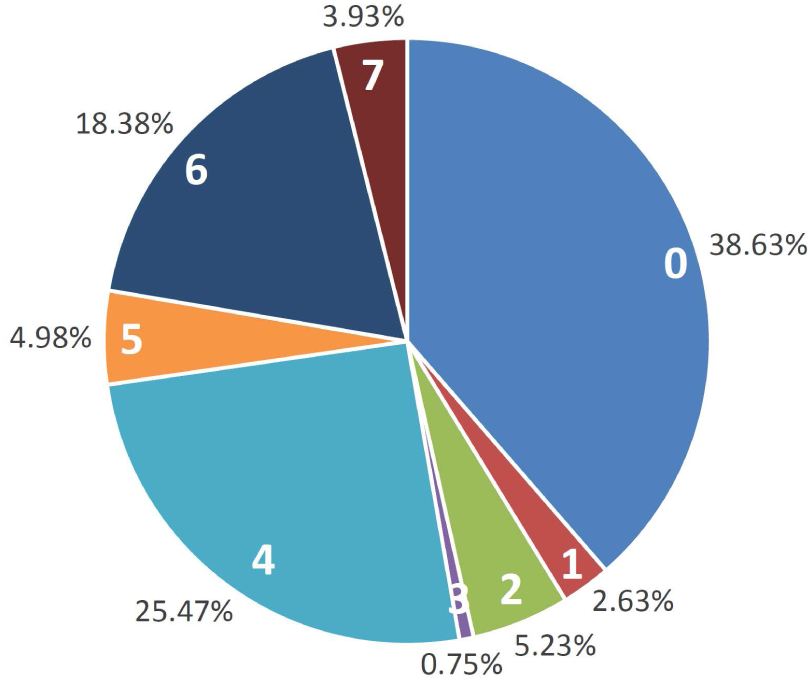
The fractions of the profile classes PC = 0 to PC =7 averaged over 35 images.

Figure 3 depicts the fractions of large cells in respect to the diagnosis. For NScHL and MCcHL, the fractions of large profiles were 17.36% and 11.06%, respectively, whereas for LA the fraction of large profiles was only 8.10%. The fractions of PC for each diagnosis are listed in Table S3 in the supplementary material. The small cell profiles built the majority of the cells with 82.64% in NScHL, 88.94% in MCcHL, and 91.90% in LA.

**Figure 3:**
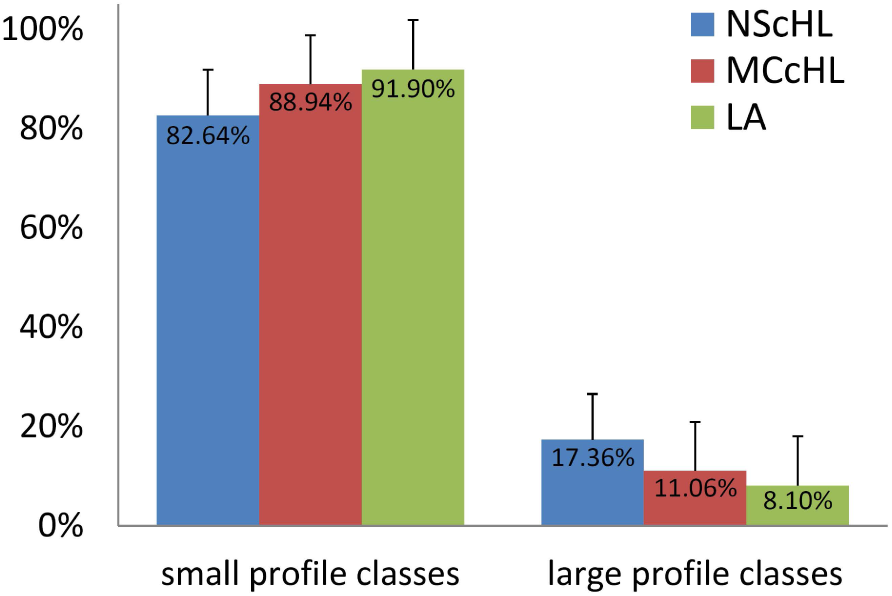
The fractions of small (PC = 0,2,4,6) and large (PC = 1, 3, 5, 7) profile classes in respect to the diagnosis. The error bars show the standard deviation. The fraction of large profiles in images of NScHL, MCcHL, and LA are statistically indistinguishable.

Because cHL is commonly characterized by the occurrence of large HRS cells, we expected to find a large number of large profiles at least in images diagnosed as cHL. Surprisingly, the measured differences in the number of large profiles in cHL and LA, respectively, were not sufficiently significant to distinguish between cHL and inflammation.

### The diameters of cells differ between cHL and LA

Cells of NScHL and MCcHL, which are most likely HRS cells, have been reported to vary between 20 *μ*m and 60 *μ*m [1, 4, 9], whereas cells of LA, which originate from activated lymphocytes, have been shown to vary between 10 *μ*m and 30 *μ*m [8, 6, 23]. Based on the distribution of the cell diameters of CD30–positive cells (Figure S4, supplementary material), we assumed the majority of cells with a diameter in the range of 12.5 to 15 *μ*m to originate either from activated lymphocytes for LA or from small HRS cells for NScHL and MCcHL. Based on the ratio of the distributions at 12.5 to 15 *μ*m, we estimated the proportion of large HRS cells to be 89.8% and 87.0% for NScHL and MCcHL, respectively. Note that, the distribution of cell diameters for LA characterizes activated lymphocytes.

To make the results more comparable with the literature values obtained by visual inspection of high resolution electron microscopy images, we used the diameter distribution of LA to subtract the background of smaller cell profiles for NScHL and MCcHL. Figure 4 depicts the corresponding corrected distributions for NScHL and MCcHL. The distributions for NScHL and MCcHL have both a maximum for diameters in the range of 22.5 and 25 *μ*m. For NScHL, the mean cell diameter was 30.6 *μ*m with a large standard deviation of 10.2 *μ*m, whereas the mean value for MCcHL was slightly smaller, 28.6 *μ*m with a standard deviation of 9.3 *μ*m. A large fraction of 9.2% and 5.6% of cells in NScHL and MCcHL, respectively, had a cell diameter larger than 50 *μ*m.

**Figure 4:**
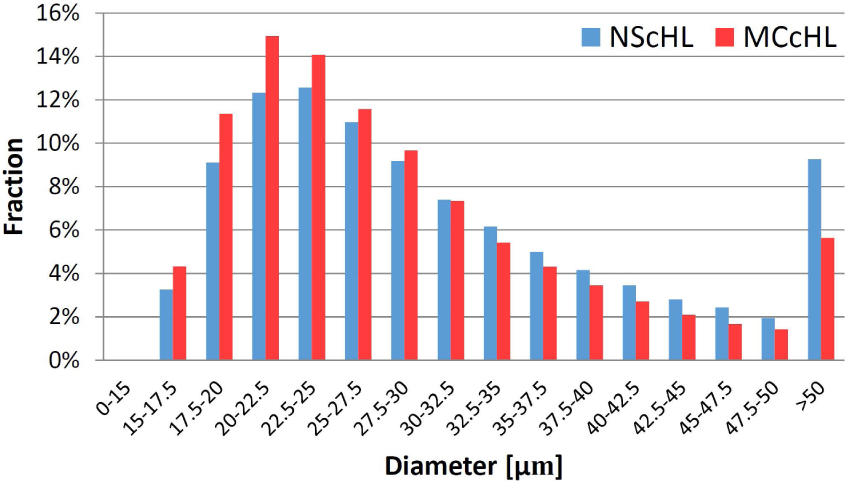
The distribution of diameters of cells. The bars show the relative fractions of of HRS cells for the diagnoses NScHL and MCcHL, respectively. The relative fractions were corrected for the background of activated lymphocytes, see text.

We computed the cell diameters defined as maximal Feret distances. The values of the maximal Feret distances may differ from the cell diameters that a pathologist determines by visual inspection. Our statistical analysis of a high number of CD30–positive cells offers a view that is complementary to a high–quality, visual inspection of individual cells. There exists no one–to–one correspondence between the computed cell diameters and those reported in the literature. Nevertheless, the values for the diameters of HRS cells, see, e.g., Figure 4, and the fraction of large HRS cells with diameters larger than 50 *μ*m were in perfect accordance with the values in the literature [1, 4, 9].

### Small, round cells prefer to stay among themselves

Figure 5 depicts the main preferences and aversions of cells in relation to their PC independent of the diagnosis. Each node represents one of the eight PC. We scored the neighborhood relations, see section *Methods*. Arrows are drawn between nodes if the absolute value of the score of neighborhood relation exceeds 50% of the maximal, possible value. Thick light grey (online version: green) and black (on-line version: red) arrows represent large positive and large negative scores, respectively. Pairs connected by light grey/green arrows favor each other as neighbors. Pairs connected by black/red arrows avoid each other as neighbors.

**Figure 5:**
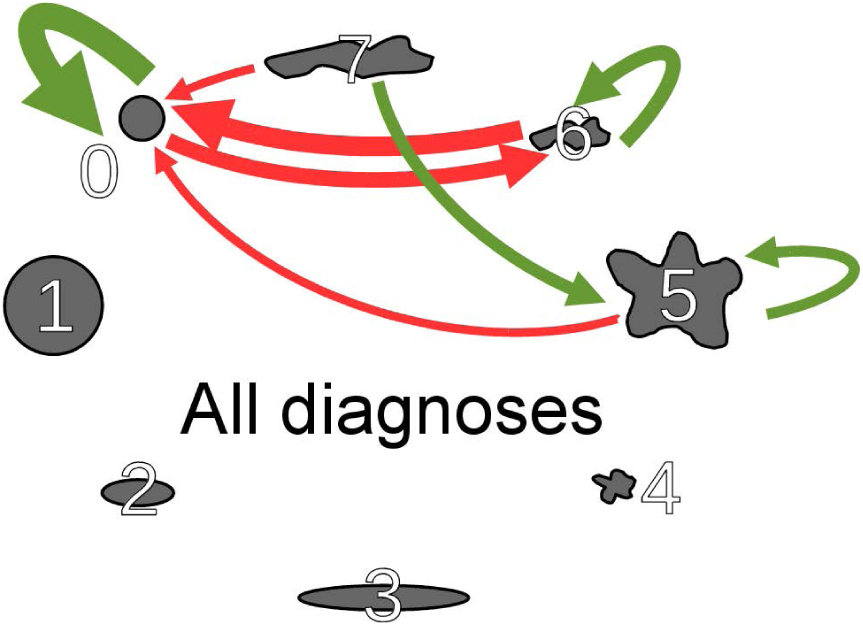
Network of neighborhood relations. Each node represents one of the eight PC. The thickness of the arrows correlates with the absolute value of the score. Light grey (online version: green) and black (online version: red) arrows represent preferences and aversions, respectively.

In Figure 5, the thick black/red arrow from PC = 6 to PC = 0 indicates a strong aversion expressed by a negative score of 74% of small, elongated, frayed profiles to have small, round cells as nearest neighbor. In 26 of the 35 images, small, round profiles were significantly underrepresented in the neighborhood of small, elongated, frayed PC. The aversion of these two PC to each other was mutual indicated by a reversed black/red arrow (from PC = 0 to class PC = 6) with a negative score of 69%. Note that, preferential and avoided neighborhood relations have not to be mutual, see Figure S5 in the supplementary material.

The light grey/green arrow from PC = 7 to PC = 5 in Figure 5 expresses the preference indicated by a positive score of 57% of large, elongated, frayed cells to have large, frayed cells in the neighborhood. Furthermore, light grey/green loop arrows show the preferences of PC = 0 (small, round), PC = 6 (small, elongated, frayed), and PC = 5 (large, frayed) toward themselves. In 89% of the images, small, round cells (PC = 0) preferred as neighbor other small, round cells (NPC = 0). The number of the combination (PC = 0, NPC = 0) was significantly high in 31 images, see Figure S3 A) in the supplementary material. In general, PC that tend to repel other classes (PC0, PC5, PC6) prefer to stay among themselves.

### Neighborhood relations differ for the three diagnoses

The three networks in Figure 6 illustrate the main preferences and aversions of cell profiles that are specific for the three diagnoses NScHL, MCcHL, and LA. In each of the three diagnoses, the small, round cells prefer to stay among themselves, exhibiting high positive scores of 83 to 92%, see Figure S3 in the supplementary material. Small, round cells are always shunned by other PC, in particular by small, elongated, frayed cells (PC = 6). Independent of the diagnoses, no PC seems to like the small, round cells, with the only exception of small, round cells themselves. Despite of common characteristics of the networks in Figure 6, significant differences in the neighborhood relation are visible for the three diagnoses.

**Figure 6:**
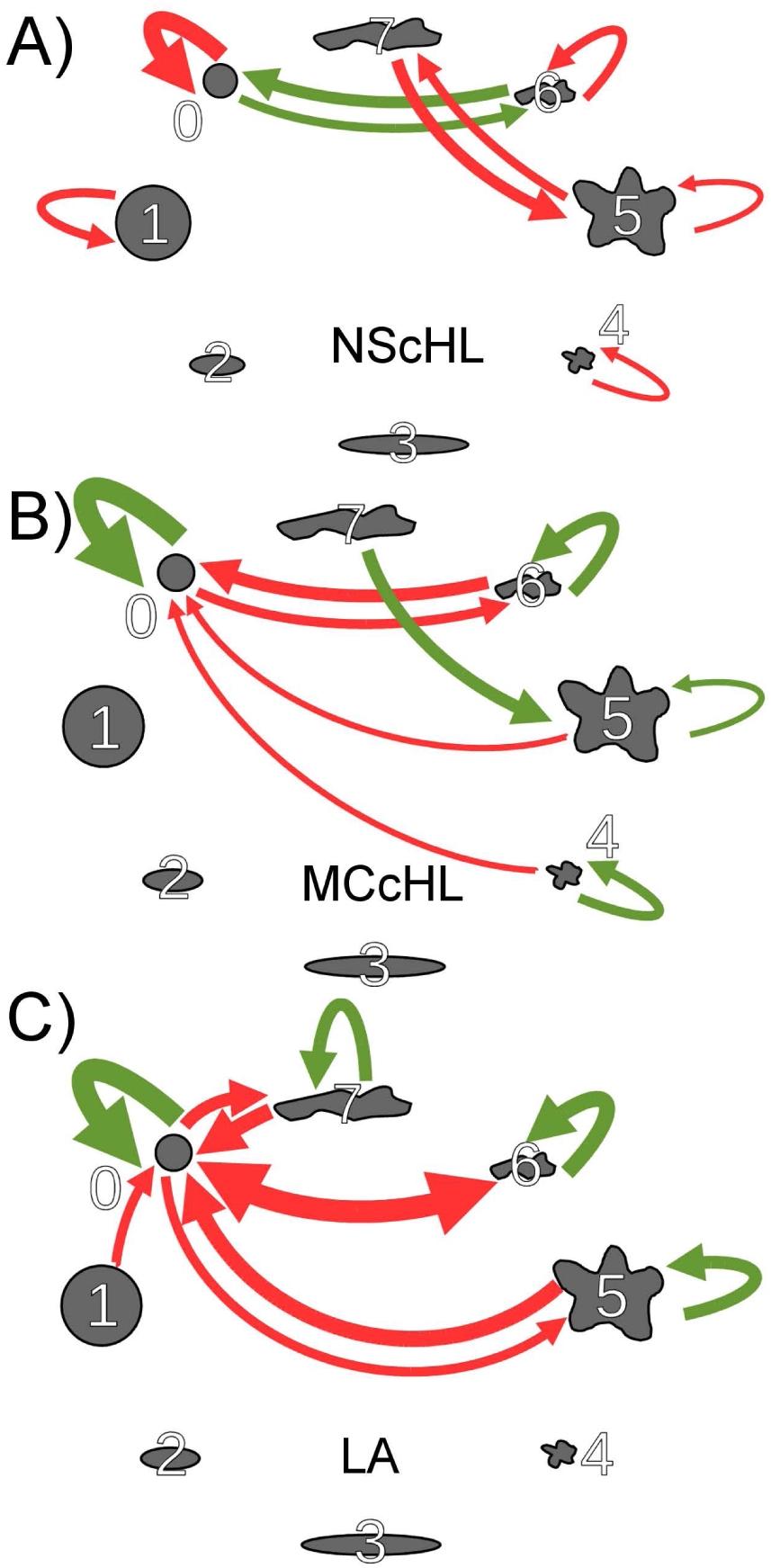
Networks of neighborhood relations A) in NScHL, B) in MCcHL, and C) in LA. Each node represents one of the eight PC. The thickness of the arrows correlates with the absolute value of the score. Light grey (online version: green) and dark grey (online version: red) arrows represent preferences and aversions, respectively.

*NScHL:* In comparison to the other diagnoses, aversions of cells to other PC is less noticeable in NScHL, see Figure 6 A). Only two black/red arrows indicate a negative score with more than 50%. Small, round cells (PC = 0) dislike small, frayed, elongated cells (PC = 6) and *vice versa*. A preferred neighborhood relation exists between large, frayed, elongated cells (PC = 7) and large, frayed cells (PC = 5).

*MCcHL::* As for NScHL, cells of PC = 7 (large, elongated, frayed) favor cells of PC = 5 (large, frayed). But, this neighbor preference becomes less pronounced in MCcHL, see Figure 6 B). Simultaneously, the aversion of cells increases to accept small, round cells, PC = 0, in their neighborhood.

*LA:* The network for LA in Figure 6 C) shows a high number of aversions between PC. Examples are the strong repulsions of PC = 0 (small, round) from PC = 5 (large, frayed), PC = 6 (small, elongated, frayed), and PC = 7 (large, elongated, frayed). Half of the profile classes (PC = 0, 5,6,7) prefer to stay among their own kind. The preferred neighborhood relation, PC = 5, NPC = 7, which is noticeable for NScHL and MCcHL is, however, absent in LA.

### Clustering of small, round cells is not caused by attraction

Table 2 shows the mean distance of a cell to its nearest neighbor. In NScHL, the cells have the smallest mean distance of 37.9 ± 6.1 *μ*m. It increases to 42.7 ± 11.9 *μ*m and 48.3 ± 18.1 *μ*m for MCcHL and LA, respectively. Within the large standard deviations, the differences in the mean distances are not statistically significant.

**Table 2:**
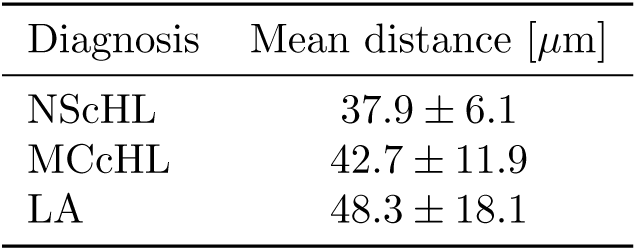
Mean distances of PC to the nearest neighbor in images of all diagnoses.

Compared to the theoretically expected distribution for spatial, randomly located cells, the distribution of distances to the nearest neighbor was shifted towards small values and had a much more narrow shape, see Figures S6, S7, and S8 in the supplementary material. The observation of a shifted and narrowed distance distribution was valid for each of the 35 images and indicates a significant clustering of the CD30–positive cells independent of the medical diagnosis. This has been verified by a graph-theoretical analysis of network properties of the corresponding cell graphs [25].

We asked whether the strong preference of small, round cells, PC = 0, to cluster together could be an effect of an attraction between these cells. For the majority of images, the ratio between the mean distances of PC = NPC = 0 nearest neighbors and the mean distances of two nearest neighbors of arbitrary classes was larger than 1, see Figure S9 in the supplementary material. The PC = NPC = 0 pairs showed large distances between them. In particular in images of low cell density, the mean distance of these pairs was up to 50% enlarged compared to arbitrary classes. This finding contradicts the hypothesis of an attractive force. The preference of these cells to cluster together must have other reasons. The preferred location of small, round cells in tissue regions that are not easily accessible by other cell types could be a possible explanation. The high motility combined with the small size of PC = 0 cells may lead to an aggressive ability to colonize healthy tissue regions with still sparse populations of CD30–positive cells.

The Pearson correlation coefficient of the mean distance between PC = NPC = 0 pairs and the log-odd value of the significance level of a preference for these pairs were positive. The correlation coefficient over all images was 0.42. The positive value of the correlation coefficient indicated that, the preference of a PC = NPC = 0–neighborhood was strongly correlated with a large distance between the cells.

The visual examination of all diagnoses showed a significantly high number of PC = NPC = 0 pairs that were located in tissue areas of low cell density, see, e.g., Figure S10 in the supplementary material. A preferential location of small, round cells in sparsely–populated tissue regions could explain the large distance between PC = NPC = 0 pairs. Interestingly, a positive correlation between large distances and neighborhood preferences occurred only for these pairs, whereas for all other combinations, the correlations were negative. Only in the cases of PC = NPC = 0 pairs, the preferences of cells of a PC to stay among themselves were strongly correlated with tendencies to keep the distances large to the next neighbors.

## 4 Conclusion

Histological images of CD30–positive cells produced in high numbers by the daily pathological praxis give a two-dimensional snapshot of the complex dynamical, three-dimensional tumor environment. The images offer a source to yield statistically significant, quantitative data for the variable, adaptive, and complex lymph node during the progression of HL. Such data are invaluable to validate biological concepts that aim to describe the lymph node on a cellular resolution and to understand diseases of the lymphatic system.

The lymph node is a structured organ with major compartments like the subcapsular sinus, B cell follicles, the T cell zone, trabecular and medullary sinuses, and blood vessels. A broad variety of cells enter the lymph node, migrate from compartment to compartment, interact with other cells, and show a complex movement in a stromal cell network [2]. Previous characterizations of the diameter of HRS cells were based on high resolution images captured by electron microscopy [8, 6]. Individual HRS cells have been identified manually by their large nuclei. The quality of electron microscopy images is excellent and identification of individual HRS cells by visual inspection is certainly still superior to any automated image analysis, but only a low number of cells has been measured. No corresponding statistical data have been presented so far. There are no studies available, which investigate the morphological properties and neighborhood relations of tumor cells in the lymph node.

The presented study provides valuable information about the morphological characteristics and neighborhood relations of cHL and LA. For the first time, a systematic analysis of a high number of 400, 000 CD30–positive cells in WSI of lymph node tissue sections of NScHL, MCcHL, and LA was performed. We provided measured values for the distances between neighbored cells, the cell diameters of HRS cells in complete lymph node sections, and morphological characteristics for each cell.

The profiles of CD30–positive cells were identified by the in-house imaging pipeline [26, 25]. Based on morphological cell features, we defined eight morphological PC to classify the cells and computed the fractions of each PC.

The distributions of the diameter of CD30–positive cells in cHL had their maxima in the range of 20 to 22.5 *μ*m For LA, the distribution had a maximum in the range of 15 to 17.5 *μ*m, which is much smaller than the estimated mean diameter of HRS cells in NScHL of 30.6 ± 10.2*μ*m The mean diameter for MCcHL was slightly smaller, about 28.6 ± 9.3*μ*m

We observed statistically significant preferences and aversions in the neighborhood relations of the PC. Round, small cells liked to stay in the neighborhood of themselves and were avoided by the other classes. The neighborhood relations between the Hodgkin cells and neighborhood relations between the lymphadenitic cells were similar, despite of some differences. For example, in LA, the cells of PC = 7 (large, elongated, frayed) were not found preferably in the neighborhood of PC = 5 (large, frayed). This lack of neighborhood preference between cells of PC = 7 and PC = 5 indicates a different spreading behavior of these cell types in tissues of cHL and lymphadenitis.

The distribution of distances between neighbored cells demonstrated a typical spatial clustering of the cells in the tissue. The only exception was a comparably large mean distance between small, round cells, which contradicts the hypotheses of an attraction that forces small, round cells to stay among themselves. Small, round cells were preferably located in regions of small cell density. Possible explanations for the overall clustering of cells are the influence of the complex structure of the lymph node and specific cell interactions caused, e.g., by chemokines or cytokines.

The development of different cell forms of HRS cells have been analyzed in cell cultures. Refusal of divided cells have been demonstrated [24]. Since no animal model of HL exists, the investigation of HL infiltration in time and space is up to now not possible. To get more insight in the dynamic cell processes, histological sections with positively immune stained tumor cells could serve as a basis for future investigations. The proliferation status of tumor cells may influence diameters and shapes of cells. We can hypothesize that small cells correspond to early stages of cell development and large cells to later stages. The forms of the cells, especially their elongation, leading to polarization, may reflect cell movement. The neighborhood analysis gives insights of their distribution at a fixed time point, including morphological cell features, and allows to draw potential strategies of tumor cell migration.

The detection of CD30–positive cells in three-dimensions by applying confocal scanning microscopy [19] will be an important aspect of experimental work. Multi-staining will allow to visualize various important players beside CD30–positive cells, as, e.g., T cells and B cells that are known to interact with malignant cells. Pattern recognition is and will further be a focus of ongoing research.

